# Phylloxera and aphids show distinct features of genome evolution despite similar reproductive modes

**DOI:** 10.1101/2023.08.28.555181

**Authors:** Zheng Li, Allen Z. Xue, Gerald P. Maeda, Yiyuan Li, Paul D. Nabity, Nancy A. Moran

## Abstract

Genomes of aphids (family Aphididae) show several unusual evolutionary patterns. In particular, within the XO sex determination system of aphids, the X chromosome exhibits a lower rate of interchromosomal rearrangements, fewer highly expressed genes, and faster evolution at nonsynonymous sites compared to the autosomes. In contrast, other hemipteran lineages have similar rates of interchromosomal rearrangement for autosomes and X chromosomes. One possible explanation for these differences is the aphid’s life cycle of cyclical parthenogenesis, where multiple asexual generations alternate with one sexual generation. If true, we should see similar features in the genomes of Phylloxeridae, an outgroup of aphids which also undergoes cyclical parthenogenesis. To investigate this, we generated a chromosome-level assembly for the grape phylloxera, an agriculturally important species of Phylloxeridae, and identified its single X chromosome. We then performed synteny analysis using the phylloxerid genome and 30 high-quality genomes of aphids and other hemipteran species. Unexpectedly, we found that the phylloxera does not share aphids’ patterns of chromosome evolution. By estimating interchromosomal rearrangement rates on an absolute time scale, we found that rates are elevated for aphid autosomes compared to their X chromosomes, but this pattern does not extend to the phylloxera branch. Potentially, the conservation of X chromosome gene content is due to selection on XO males that appear in the sexual generation. We also examined gene duplication patterns across Hemiptera and uncovered horizontal gene transfer events contributing to phylloxera evolution.

## Introduction

Aphids (Insecta: Hemiptera: Aphididae) are a monophyletic group of about 5,000 species that feed on plant sap and include some globally distributed agricultural pests. They show remarkable life cycles incorporating cyclical parthenogenesis, in which several asexual, all-female generations are interspersed with a single sexual generation. Cyclical parthenogenesis depends on modifications of meiosis, including non-meiotic reproduction during all-female generations, elimination of an X chromosome to produce XO sons, and production of only X sperm by males to yield only XX daughters (Moran 1992; Davis 2012, Blackman 1980).

Aphids also exhibit distinctive patterns of genome evolution. Recently, chromosome-level assemblies have revealed that aphid X chromosomes display long-term conservation of gene content and arrangement, contrasting with relatively frequent autosomal rearrangements including autosomal translocations (Li et al. 2019; Mathers et al. 2021). Despite this conservation of gene content, aphid X-linked genes show low expression and elevated rates of nonsynonymous substitution, consistent with weak purifying selection, and a trend towards male-biased expression (Jaquiéry et al. 2018; Li et al. 2019; Jaquiéry et al. 2013; Mathers et al. 2021). Genes that are highly expressed, critical to all life stages (for example, those encoding ribosomal proteins), and/or single-copy are heavily concentrated on autosomes and largely lacking in the X chromosome. These observations suggest that selection favors placing functionally critical genes on autosomes, and that genes remaining on the X chromosome undergo less stringent purifying selection.

The cyclical parthenogenesis of aphids is hypothesized to underlie this distinctive pattern of genome evolution (Jaquiéry et al. 2018; Li et al. 2019). If true, these genomic features are predicted to extend to the phylloxera (family Phylloxeridae), a related lineage in the same infraorder as aphids, Aphidomorpha. Aphids and phylloxera diverged over 160 MYA (Ren et al. 2013). These two lineages share features reflecting their shared ancestry, most notably cyclical parthenogenesis and XO sex determination (Morgan 1909; Forneck and Huber 2009). There are also several key differences. One is that, during asexual reproduction, aphids are viviparous, while phylloxera produce eggs. A second difference is that aphids contain intracellular bacterial endosymbionts that provide amino acids (Shigenobu et al. 2000; Chong et al. 2019) and support these symbionts with genes acquired by horizontal gene transfer (HGT) from bacteria (Nakabachi et al. 2014; Smith et al. 2022), while phylloxera lack endosymbionts and the corresponding HGT genes (Rispe et al. 2020).

The grape phylloxera, *Daktulosphaira vitifoliae*, has historically had a global impact on the grape and wine industries. A recent study characterized its genome and elucidated its population and global invasion history (Rispe et al. 2020). However, this assembly is highly fragmented, preventing chromosome-level comparisons between aphids and phylloxera.

In this study, we used Dovetail Omni-C technologies to assemble a chromosome-level genome for the grape phylloxera. Using analyses of genomes of 30 other hemipteran species with standard sexual life cycles and XO/XY sex determination, we tested whether the distinctive features of aphid genome evolution originated with the origin of cyclical parthenogenesis, and thus extend to phylloxera but not to other hemipterans. In addition, we used the new genome assembly to examine the extent and pattern of gene duplication in phylloxera as compared to aphids and to identify HGT events that have contributed to phylloxera evolution.

## Results

### Genome assembly and annotation of grape phylloxera

The assembled genome was produced using data from a proximity ligation protocol (Dovetail Omni-C) incorporating 11.6 Gb of new Illumina reads and the previously published draft genome assembly (Rispe et al. 2020). Our genome has a total length of 282.86 Mb and an N50 of 45.89 Mb (supplementary table S1). Five scaffolds are larger than 30 Mb, suggesting a haploid chromosome count of n = 5 (supplementary fig. S1). The total length of the chromosome-level scaffolds is 259.64 Mb, which is 91.8% of the total length of the assembly.

We used BUSCO (Benchmarking Universal Single-Copy Orthologs) to evaluate the completeness of our genome assembly. Querying the single copy orthologs of Hemiptera resulted in a BUSCO score for the genome assembly of 99.5% (99.1% single and duplicated, 0.4% fragmented, 0.5% missing). The BUSCO score for the five chromosome-level scaffolds alone was 99.1% (98.7% single and duplicated, 0.4% fragmented, 0.9% missing).

The NCBI RefSeq annotation pipeline was used to annotate the genome (O’Leary et al. 2016). We used WindowMasker (Morgulis et al. 2006) to mask repetitive elements, which made up 53.26% of the genome. We then aligned 18 phylloxera transcriptomes containing 1,698,647,388 reads onto the repeat-masked genome. We predicted a total number of 17,104 annotated genes and pseudogenes, with 14,650 protein-coding genes. Overall, 15,582 predicted genes were annotated on the five chromosome-level scaffolds, with 2,670 on the X chromosome and 12,912 genes on the four autosomes (Table 1).

**Table 1.** Summary of the chromosome-level assembly of the grape phylloxera genome.

| **Category** | **Length (Mb)** | **No. of genes** |
| --- | --- | --- |
| Chromosome 1 | 89.58 | 5,226 |
| Chromosome 2 | 45.89 | 2,936 |
| Chromosome 3 | 43.29 | 2,712 |
| Chromosome 4 | 33.59 | 2,038 |
| Chromosome X | 47.28 | 2,670 |
| All Chromosomes | 259.64 | 15,582 |
| Total Assembly | 282.86 | 16,912 |

### Confirmation of X chromosome

To identify the X chromosome in the phylloxera genome, we first looked at genome synteny between the grape phylloxera and the pea aphid (*Acyrthosiphon pisum)* (supplementary fig. S2). The second-largest chromosome (47.28 Mb) showed extensive gene synteny with the pea aphid X chromosome. To confirm the identity of the X chromosome in phylloxera, we obtained Illumina reads from sexual males and from females and mapped them to our five chromosome-level scaffolds. The second-largest scaffold had about half of the normalized sequencing read depth ratio for males when compared to other chromosomes (supplementary fig. S3), confirming that it is the X chromosome. We named the four autosomes as chromosomes 1-4, ordered from longest to shortest (Table 1).

### Synteny evolution of grape phylloxera and other hemipteran insects

Synteny between species was generated by MCScanX (Wang et al. 2012) and was visualized with SynVisio (https://github.com/kiranbandi/synvisio). Based on comparisons of assemblies for different species, we observed numerous rearrangements and shuffling of syntenic regions among aphid autosomes. In contrast, gene content and synteny of the X chromosome were highly conserved between phylloxera and aphids (Fig. 1). Most Aphidomorpha, including phylloxera, possess a single conserved X chromosome. In the *Hormaphis* (Hormaphidinae) and *Eriosoma* (Eriosomatinae) (Fig. 1), the X is split into two chromosomes but these retain conserved syntenic regions corresponding to the X.

**Fig. 1.**
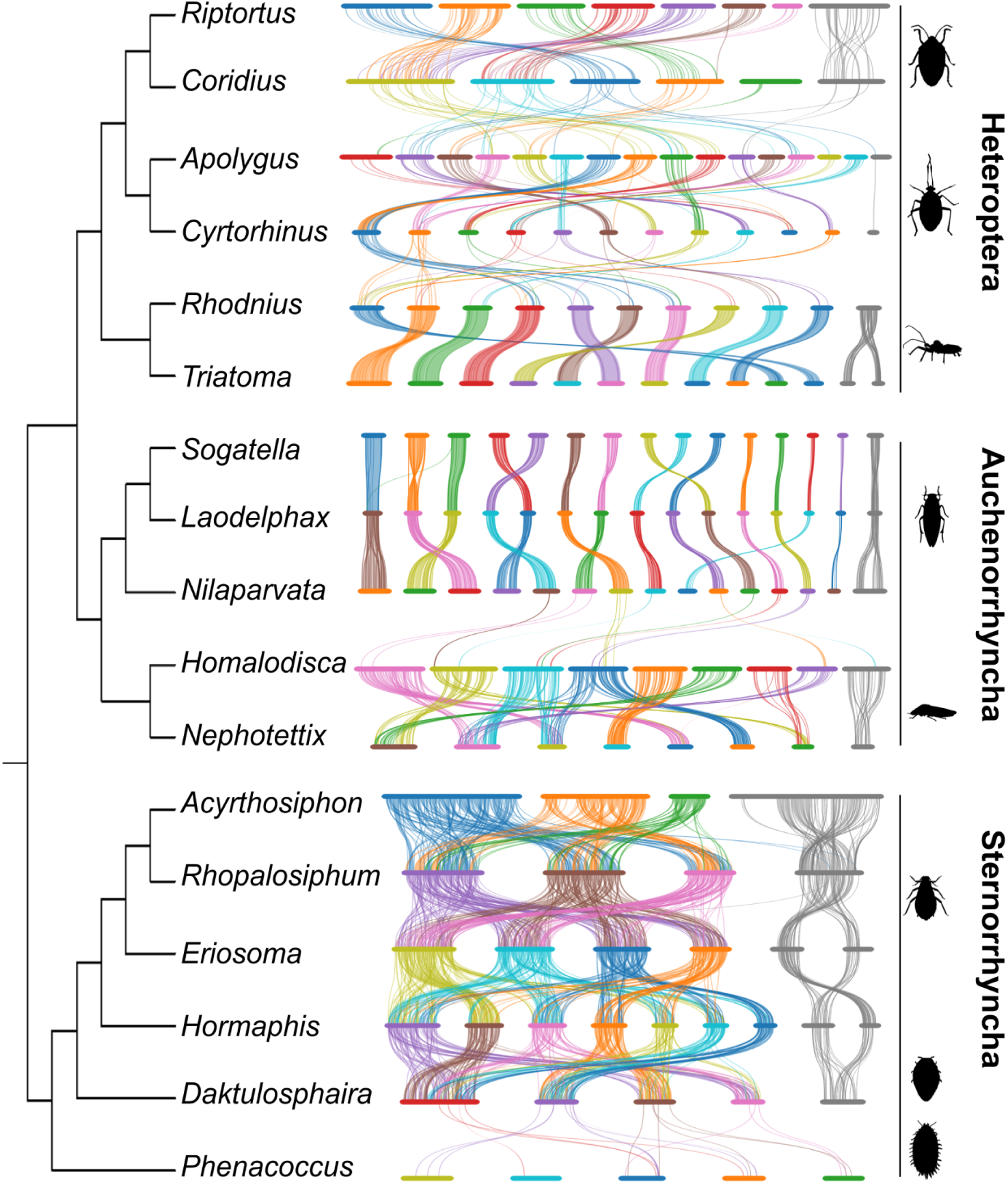
Pairwise syntenies (gene order) of Heteroptera (true bugs), and Auchenorrhyncha (planthoppers, leafhoppers, and relatives), and Sternorrhyncha (aphids and relatives). Bars represent chromosomes. The X chromosome (s) for each species is colored gray and located on the far right. Autosomes are in other colors and ordered by size from large to small (left to right). The length of the bars is proportional to the length of the chromosome-level scaffolds in the assemblies. The cladogram is based on Johnson et al. 2018.

To study whether elevated autosomal rearrangements are shared by Aphidomorpha and are associated with the origin of cyclical parthenogenesis, we tested if this unusual pattern of chromosome evolution extended to phylloxera. We found a high level of autosomal rearrangements when comparing aphids to phylloxera. However, this elevated rate of autosomal rearrangements could be unique to the aphid branch. To determine whether the difference between autosomes and X chromosome extends to phylloxera, we separately compared aphids and phylloxera with *Adelges cooleyi*, a member of the family Adelgidae, a third related lineage undergoing cyclical parthenogenesis. Thus, if parthenogenesis is responsible for autosomal rearrangements, the elevated autosomal rearrangements should be present in both comparisons.

However, the elevation was only found when comparing aphids to the other two lineages. In contrast, the comparison of phylloxera and adelgids showed strong synteny conservation for autosomes, suggesting that elevated autosomal rearrangements are restricted to Aphididae.

To further study whether Aphididae uniquely display conservation of gene content of the X chromosome contrasting with a higher rate of interautosomal translocations, we extended our analyses to the other two major hemipteran clades, Heteroptera and Auchenorrhyncha, for which sufficient chromosome-level assemblies were available (supplementary table S2). We found no evident differences in the frequencies of interchromosomal rearrangements between autosomes and X chromosomes in these two clades (Fig. 1).

To quantify the difference in chromosomal evolution between Aphididae and other hemipteran lineages, we estimated approximate rates of interchromosomal rearrangement events, as calibrated using estimated divergence dates of the included species. These events include translocations and fusions/fissions; however, most of them are translocations. Because the signature of small chromosomal rearrangements can be ambiguous and difficult to infer, we focused on major rearrangements between species pairs with a minimum 50 MY of divergence (supplementary table S3, S4). We classified chromosomes into four categories: Aphididae X chromosomes, Aphididae autosomes, other hemipteran X chromosomes, and other hemipteran autosomes. Aphididae autosomes have a mean of 0.025 major translocation events per MY. Averages for the other three categories range from 0.003 to 0.014 events per MY. Overall, Aphididae autosomes have a significantly higher rate of major rearrangements compared to other hemipteran autosomes (Mann–Whitney *U* = 2851.5, p *<* 10^−15^) and X chromosomes (Mann–Whitney *U* = 299.5, p *<* 10^−4^). Aphididae X have a significantly higher rate of major rearrangements compared to other hemipteran autosomes (Mann–Whitney *U* = 734, p *<* 0.004), and rates of rearrangement are not significantly different between X chromosomes and autosomes for other hemipterans (Mann–Whitney *U* = 703.5, p = 0.82) (Fig. 2, supplementary table S5).

**Fig. 2.**
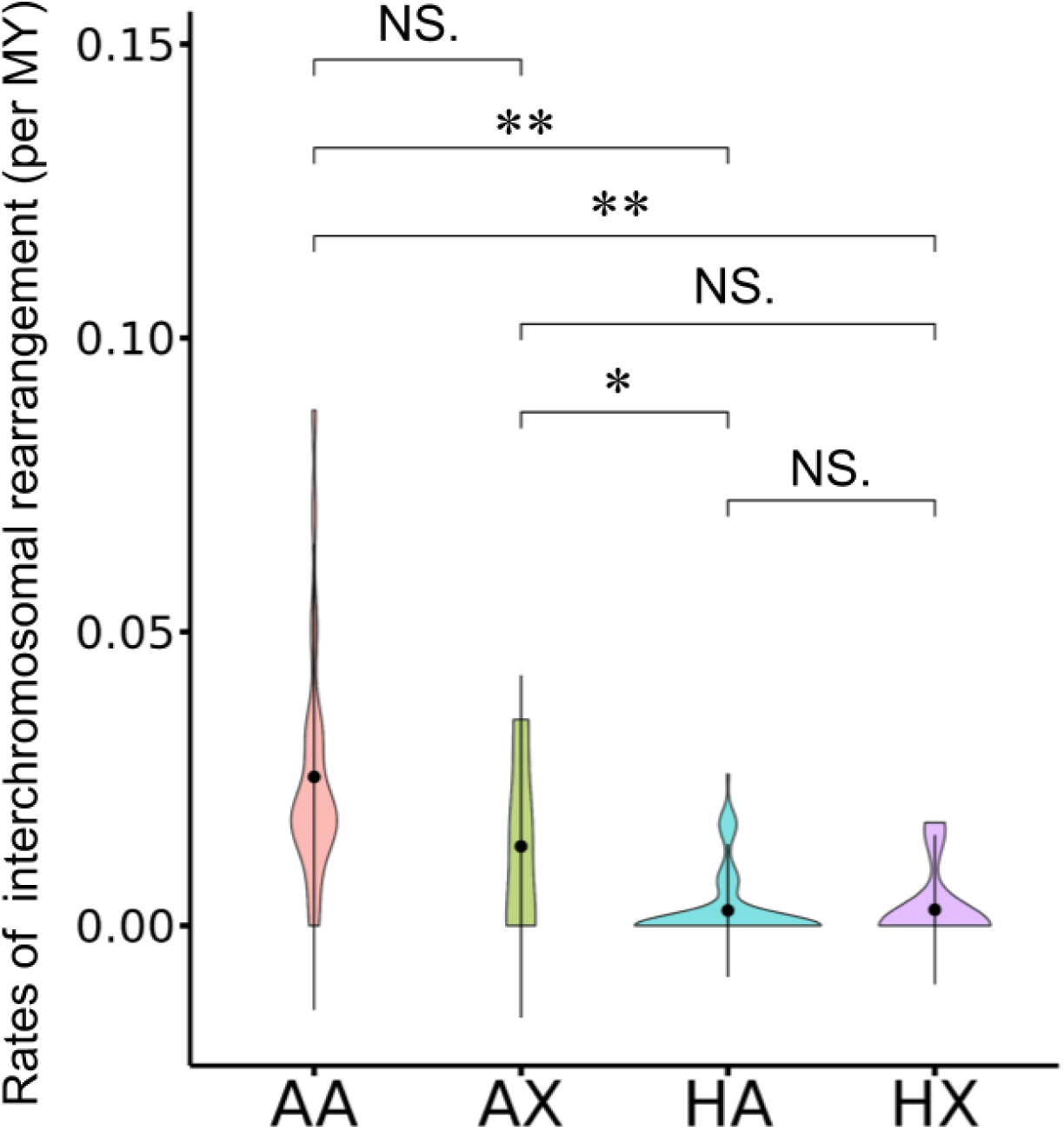
The rates of interchromosomal rearrangement of Aphididae and other hemipteran chromosomes (AA, Aphididae autosomes; AX, Aphididae X chromosomes; HA, other hemipteran autosomes; HX, other hemipteran X chromosomes). The rate of interchromosomal rearrangement within each category of chromosome is shown in the violin plots. The black dot indicates the mean, the thick black bars represent the standard deviation of the data, and the color shading represents the density of data points. The comparisons of the Aphididae autosomes to other categories by the two-sample Mann-Whitney U test are provided. ‘**’ represents Bonferroni-corrected *p* < 0.002, ‘*’ represents Bonferroni-corrected *p* < 0.01

### Sequence evolution of aphids and phylloxera

Compared to genes on autosomes, X-linked genes in aphids show a faster rate of gene sequence evolution, particularly at nonsynonymous sites (Jaquiéry et al. 2012, 2013; Purandare et al. 2014; Jaquiéry et al. 2018; Li et al. 2020). To test if this faster-X pattern is shared by phylloxera, we estimated sequence divergence between phylloxera and pea aphid and between phylloxera and adelgid for gene pairs or orthologs within syntenic blocks (supplementary table S6). Between phylloxera and pea aphid, we found that the mean divergence at nonsynonymous sites is higher for X-linked genes (mean d*N*_X_ = 0.30) than for genes on autosomes (mean d*N*_A_ = 0.23, Mann–Whitney *U* = 296745.5, p *<* 10^−5^). The divergence at synonymous sites is similar for genes on the X chromosome and on autosomes (mean d*S*_X_ = 1.20, mean d*S*_A_ = 1.25, Mann–Whitney *U* = 393876.5, p *<* 0.04). The d*N*/d*S* ratio is higher for genes on the X compared to genes on autosomes (mean d*N*_X_/d*S*_X_ = 0.30, mean d*N*_A_/d*S*_A_ = 0.21, Mann–Whitney *U* = 297275.5, p *<* 10^−5^). Similarly, between phylloxera and adelgid, the mean divergence at nonsynonymous sites and d*N*/d*S* ratio are higher for X-linked genes (mean d*N*_X_ = 0.20, mean d*N*_X_/d*S*_X_ = 0.17) than for genes on autosomes (mean d*N*_A_ = 0.15, Mann–Whitney *U* = 657048, p *<* 10^−10^, mean d*N*_A_/d*S*_A_ = 0.14, Mann–Whitney *U* = 685454, p *<* 10^−7^) (supplementary table S6).

### Gene duplication in phylloxera

Based on recent genomic studies, aphids genomes show high but variable rates of gene duplication (Julca et al. 2020; Mathers et al. 2017; Li et al. 2019; Fernández et al. 2020) and ancient large-scale gene duplications (Julca et al. 2020). This variation has resulted in wide differences in the total number of protein-coding genes, ranging from 14,089 to 31,435 genes (supplementary table S7). In the grape phylloxera genome, we identified 9,137 genes with duplicates, around 62% of total genes (supplementary fig. S4, supplementary table S7), a value near the lower end of the range for Aphidomorpha overall (54-81% of total genes). Relatively low numbers of gene duplications are also observed in Hormaphidinae and *Schlechtendalia* compared to many Aphidinae.

By using synteny and gene locations on chromosome-level assemblies, we further classified gene duplications into four modes: tandem (immediately adjacent), proximal (within 20 flanking genes on a chromosome), dispersed (not within 20 flanking genes), and segmental duplications (anchored within colinear gene blocks) (supplementary fig. S4). We found little evidence of segmental or other large-scale gene duplications. The majority of gene duplications in aphids and phylloxera are dispersed. However, this may reflect the long evolutionary time periods involved, during which larger duplication events are broken up by chromosome rearrangements in autosomes.

### HGT in phylloxera

We used protein annotation and the Alienness pipeline (Rancurel et al. 2017) to look for HGT genes in the phylloxera genome. As predicted based on the lack of endosymbionts in phylloxera, we could not detect homologs of the HGT genes of aphids that are hypothesized to interact with the *Buchnera* endosymbionts (Moran et al. 2009). In addition to a previously documented HGT of two genes underlying carotenoid biosynthesis (Rispe et al. 2020), we identified four genes as putative HGT genes: two belong to glycoside hydrolase family 32 and two are identified as plasmid pRiAb ORF-3 protein-like genes (plasmid ORF genes) (supplementary table S8, S9). These HGT genes have amino acid sequence similarities of <60% to the closest bacterial sequences. Both glycoside hydrolase genes and one of the plasmid ORF genes carry an intron, indicating that they do not result from bacterial contamination of the DNA sample. We also used PCR amplification with DNA from phylloxera from different sampling locations to further confirm that all HGT candidates are from the grape phylloxera genome (supplementary fig. S5, supplementary table S10). Based on our genome assembly, the two glycoside hydrolase genes are found at nearby locations on Chromosome 3, while the two plasmid ORF genes are located on the X chromosome.

To access phylogenetic placement of these HGT events, we assembled a dataset with transcriptomes of Adelgidae and Phylloxeridae and genomes of the grape phylloxera and other aphids. We inferred gene presence and absence of these putative HGT genes across Aphidomorpha. Glycoside hydrolase genes appear to have been acquired in a shared ancestor of Phylloxeridae after its split from Aphididae and Adelgidae, as these genes are also found in one unidentified *Phylloxera* species and in *Phylloxerina nyssae*, which represents the deepest branching lineage in the Phylloxeridae (supplementary table S11). Similarly, the plasmid ORF genes are found in all Phylloxeridae e but not in the Adelgidae or Aphididae.

To study expression patterns of the HGT genes, we mapped RNAseq data from three major life stages of the grape phylloxera (egg containing embryo, leaf gall, and root gall) to our genome assembly (supplementary table S9). All four putative HGT genes are expressed at detectable levels in each stage. Both glycoside hydrolase genes are expressed significantly more highly in the root gall stage compared to leaf gall stage, while both plasmid ORF genes are expressed, but not at significantly different levels, in any stage (supplementary table S9).

## Discussion

Life cycles that incorporate unusual reproductive modes present an opportunity to understand the forces acting on genome evolution, including those acting on sex chromosomes. The recent availability of many more chromosome-level genome assemblies can illuminate how these forces may vary due to life cycle variation.

In this study, we used a new chromosome-level assembly for the grape phylloxera to determine whether the unusual life cycle shared by aphids and phylloxera has affected their genome evolution similarly. Our new assembly is significantly improved compared to the previous phylloxera genome assembly (Rispe et al. 2020). Our haploid chromosome count agrees with previous karyotyping studies (Maillet 1957; Forneck et al. 1999). Overall, the agreement between our assembly and flow cytometry estimates of genome size and karyotyping supports the high quality and accuracy of this genome assembly.

### Elevated rates of interautosomal translocations in Aphididae

Our findings are consistent with previous results in aphids, showing long-term conservation of gene content despite elevated gene sequence evolution for the X chromosome, contrasting with frequent interautosomal translocations (Li et al. 2019; Mathers et al. 2021). One hypothesis for the high rate of interautosomal translocations is their accumulation during aphids’ asexual generations. Other hemipterans have fewer annual generations, and all generations are sexual, likely constraining opportunities for translocations. Substantial evidence supports high rates of translocations during asexual aphid generations. Karyotypes in aphids that are exclusively asexual evolve rapidly (Blackman 1980), and translocations and fusions occur during the short-term evolution of asexual pest aphid species (Monti et al. 2012b) and in laboratory cultures (Spence and Blackman 2000). In predominantly asexual aphids, exemplified by species of the tribe Tramini, karyotypes are highly variable (Blackman et al. 2000; Normark 1999). However, by comparing synteny between aphids, phylloxera, and adelgids, we found these features of chromosome evolution have uniquely characterized Aphididae and not shared with Phylloxeridae and Adelgidae. Our findings reject the hypothesis that cyclical parthenogenesis itself can explain the pattern of chromosome evolution in aphids.

An alternative explanation for the high rate of autosomal rearrangements is the presence of holocentric chromosomes in aphids (Schubert and Lysak 2011; Mandrioli and Manicardi 2020). In plants, holocentric chromosomes can facilitate rearrangements, yielding variable karyotypes (Hofstatter et al. 2022), and elevated chromosomal evolution has been observed in some other groups with holocentric chromosomes (Höök et al. 2023; Ruckman et al. 2020; Fradin et al. 2017). However, in aphids, a high rate of rearrangements is likely not driven by holocentricity. All other hemipteran insects have holocentric chromosomes (Tree of Sex Consortium 2014), yet they have much lower rates of translocations than aphids. Indeed, most hemipterans show lower rates of chromosome evolution than most other insect orders (Ruckman et al. 2020; Kuznetsova et al. 2021).

### Conservation of X chromosome gene content in Aphidomorpha

Most likely, aphids undergo relatively frequent rearrangements in all chromosomes during asexual generations, but the sexual generation imposes a stronger selective sieve on the X than on autosomes. For example, in asexual populations of *Myzus persicae*, chromosomal translocations involve both autosomes and the X chromosome (Monti et al. 2012a). Furthermore, the X chromosome does undergo intrachromosomal rearrangements (Fig. 1), indicating that it is not immune to recombination events that affect large-scale gene synteny.

Translocations involving autosomes or the X chromosome will often have little fitness consequence during asexual female generations, which are freed from constraints of homolog pairing during meiosis. However, translocations affecting gene content of the X are expected to be highly negatively selected during the sexual generation, as XO males would lack essential genes, or suffer deleterious changes in gene dosage as proposed previously (Li et al. 2019; Mathers et al. 2021). An illustration of this kind of selection can be seen for the ribosomal RNA (rRNA) genes, which are located on the X chromosome. The rRNA locus can be eliminated from one homolog through mitotic recombination during asexual generations in laboratory culture (Blackman and Spence 1996). Clearly, this deletion would be lethal in males. Thus, selection on males may explain the lack of translocations affecting the X chromosome.

Despite the general conservation of X chromosome gene content, we note that the rRNA genes appear not to be confined to the X chromosome in phylloxera or adelgids, as they are in aphids. Of the 35 rRNA operons identified in the grape phylloxera, only two were on the X chromosome, 31 were on autosomes, and 2 were on short contigs not placed on chromosomes.

Potentially, some other feature of the unusual meiosis of aphids explains the conservation of X chromosome gene content. Chromosomal behavior during meiosis differs even among aphid species, and how chromosome inheritance mechanisms affect evolution of chromosome gene content is not clear (Manicardi et al. 2015).

A recent study proposed that gene dosage for X-autosome fusions under somatic X chromosome loss might explain the conservation of aphid X chromosomes (Roy 2021). However, this hypothesis cannot explain chromosome evolution in aphids generally, as it assumes intensive inbreeding among aphids. Many aphids outbreed, and outbreeding is enforced by life cycles in many aphid groups in which males and females fly separately to alternative host plants (Moran 1988; Hardy et al. 2015).

### Sequence evolution of aphids and phylloxera

The faster-X gene sequence evolution observed in our study and in previous investigations (Jaquiéry et al. 2012, 2013; Purandare et al. 2014; Jaquiéry et al. 2018; Li et al. 2020) has been attributed to the lower expression of X-linked genes and the tendency of these genes to be male-biased, as selection on males is infrequent during the life cycle. The aphid X chromosome is highly methylated and transcribed at a lower rate than autosomes (Mathers et al. 2019), a finding that is consistent with the explanation of relaxed purifying selection on X-linked genes. Given that XO sex determination and similar X chromosome gene content, the faster-X pattern was predicted to extend to phylloxera. Our findings confirmed that X-linked genes in phylloxera have higher rates of amino acid replacement, despite similar rates of silent substitution, as compared to genes on autosomes.

### HGT in the phylloxera

Recent genomic surveys show that HGTs are widespread in insects (Li et al. 2022a), including hemipteran species (Nováková and Moran 2012; Li et al. 2022b; Mao et al. 2018). We found at least two previously undocumented HGTs with bacterial origins. These include two nearby glycoside hydrolase genes that contain an intron at a similar position. These likely reflect HGT to an ancestor of Phylloxeridae, followed by intron gain, then followed by tandem duplication. Potentially, these genes are used for general enhancement of herbivory, where they contribute to breakdown of plant polysaccharides. Previous studies have found glycoside hydrolase genes that were horizontally transferred from different bacterial sources into herbivorous insects in several orders, including leafhoppers, several beetle lineages, and silkworms (Li et al. 2022a; McKenna et al. 2019; So et al. 2022; Pauchet and Heckel 2013; Miyazaki et al. 2020; Mao et al. 2018; Li et al. 2022b). The other HGT genes are homologous to uncharacterized ORFs on bacterial plasmids and are expressed at three major life stages of the phylloxera, where their functions are unclear. Based on a recent survey of HGT in insects, genes homologous to genes on bacterial plasmids have been transferred to different insect lineages at least three times (Li et al. 2022a).

## Conclusion

An implication from our study is that the accelerated rate of autosomal rearrangements in aphids cannot be explained by their cyclically parthenogenetic life cycle. To explain the X chromosome conservation in aphids and phylloxera, we hypothesize that the absence of this elevated translocation rate on the X chromosome reflects purifying selection on males during the annual sexual generation. One prediction is that the aphid X will exhibit more translocations to and from autosomes in lineages of Aphidomorpha that have eliminated the sexual phase. Our study raises more questions: What mechanism drives elevated rates of interautosomal rearrangements in Aphididae? Do higher rates of interautosomal rearrangements in aphids lead to higher speciation rates compared to phylloxera and adelgids?

## Materials and Methods

### Sample preparation for genome sequencing

Multiple *D. vitifoliae* Pcf7 clone females were collected from two individual plants (two different rootstocks of grapes) from the Bordeaux collection. For a high-quality genome assembly, a total of 200 mg of fresh material from 1,500 leaf-galling female phylloxera was frozen and shipped to Dovetail Genomics (Santa Cruz, CA, USA). All individuals were used for DNA extraction and Hi-C library preparation. The library was sequenced on an Illumina HiSeq X platform to produce approximately 30x sequence coverage.

### Assembly of the grape phylloxera genome

To assemble the *D. vitifoliae* genome, published *de novo* draft genome assembly (Rispe et al. 2020) and 11.6 Gb Dovetail proximity ligation reads were used as input data for HiRise genome scaffolding. The detailed assembly method for the draft genome can be found in Rispe et al. 2020. HiRise assembler version v2.1.6-072ca03871cc was used for scaffolding with default parameters (Putnam et al. 2016). Proximity ligation library sequences were aligned to the draft input assembly using bwa (Li and Durbin 2009) with defaults. The separations of proximity ligation library read pairs mapped within draft genome scaffolds were analyzed by HiRise to produce a likelihood model for the genomic distance between read pairs, and the model was used to identify and break putative misjoins, score prospective joins, and make joins above a threshold (https://omni-c.readthedocs.io/en/latest/). Overall, 1,850 joins were and 30 breaks were made to the input assembly. To evaluate the completeness of our genome assembly, BUSCO version 3.0.2 (Simão et al. 2015) was used on the chromosome-level assembly with the single-copy orthologous gene set for Hemiptera from OrthoDB version 9 (Zdobnov et al. 2017).

### Genome annotation

The NCBI Eukaryotic Genome Annotation Pipeline was used for genome annotation (O’Leary et al. 2016). Repeat families found in the genome assemblies of *Daktulosphaira vitifoliae* were identified and masked using WindowMasker (Morgulis et al. 2006). Over 20,000 transcripts of phylloxera and high-quality proteins of phylloxera and other closely related insects were retrieved from Entrez, aligned to the genome by Splign (Kapustin et al. 2008), Minimap2 (Li 2018), or ProSplign (https://www.ncbi.nlm.nih.gov/sutils/static/prosplign/prosplign.html). Additionally, 1,698,647,388 reads from 18 phylloxera RNA-Seq datasets were also aligned to the repeat-masked genome. Protein, transcript, and RNA-Seq read alignments were passed to Gnomon for gene prediction. The final annotation set was assigned to models based on known and curated RefSeq and models based on Gnomon predictions. The overall quality of the annotations was assessed using BUSCO v4 (Seppey et al. 2019). The detailed annotation pipeline can be found at https://www.ncbi.nlm.nih.gov/genome/annotation_euk/process/.

### Assignment of the X chromosome and autosomes

The X chromosome was assigned following the method previously used in the pea aphid and psyllid genomes (Li et al. 2019, 2020). We mapped whole genome sequencing reads from male and asexual female individuals back to our chromosome-level genome assembly. The male sequencing reads were generated through this study (BioProject: PRJNA929591, Accession: SRR23285932). The asexual female sequencing reads were obtained from the previous phylloxera genome project (Rispe et al. 2020) through GenBank (BioProject: PRJNA588186, Accession: SRR10412121). The sequencing reads were cleaned with Trimmomatic version 0.38 (Bolger et al. 2014). The clean reads were mapped to the chromosome-level assembly using Bowtie2 version 2.3.4.3 (Langmead and Salzberg 2012) with default parameters. The resulting SAM files were converted to BAM files, sorted, and indexed using SAMtools version 1.9 (Danecek et al. 2021). We estimated the sequencing depth based on 10-kb sliding windows with 2-kb steps, and the sequencing depth of each window was estimated using Mosdepth version 0.2.3 (Pedersen and Quinlan 2018). We normalized the overall sequencing depths among male individuals and female individuals based on methods used in Li et al. 2020. The overall sequencing depth distribution was plotted using a violin plot in ggplot2 version 3.2.1 (Wickham 2016). The X chromosome was assigned to the chromosome that had about half the ratio of sequencing depth between males and females compared to the others.

### Synteny analyses of Hemipteran genomes

We used MCScanX (Wang et al. 2012) to evaluate whole genome synteny between hemipteran species. Eight aphid genomes were downloaded from Aphidinae comparative genomics resource on Zenodo (https://zenodo.org/record/5908005#.Y255M3bMI5Y); other hemipteran genomes were obtained from published datasets (supplementary table S2). All vs. all blastp searches with genome protein sequences were performed with an *e*-value of 1e-10. MCScanX was used to generate synteny data between species with defaults. SynVisio (https://github.com/kiranbandi/synvisio) was used to display syntenies. As genomes of multiple species are available for some aphid clades, we selected *Acyrthosiphon pisum* for Macrosiphini, *Rhopalosiphum maidis* for Aphidini, and *Eriosoma lanigerum* for Eriosomatinae (Figure 1).

The level of interchromosomal rearrangements between chromosomes was quantified with synteny comparisons between aphids and between other hemipteran insects. A total of 5 aphid and 11 other hemipteran genome assemblies were used. For each species, we selected a genome from the sister tribe or family in the dataset for pairwise synteny analysis with MCScanX. All selected pairs had a minimum of 50 million years of divergence (supplementary table S3). All chromosomes were classified into four categories: aphid autosomes (*n=*27), aphid X chromosomes (*n=*9), other hemipteran autosomes (*n=*114), and other hemipteran X chromosomes (*n=*12) (supplementary table S4). Identification of the X chromosome was based on genomic confirmation from previous studies or sequence homology with a confirmed X chromosome. For each chromosome, we recorded the number of translocations between chromosomes of the selected species pairs using data generated from MCScanX. To estimate the approximate rates of chromosomal rearrangement events without reconstructing ancestral chromosomes, we divided the number of chromosomal rearrangement events by the estimated divergence dates of the species pairs. We then tested if the rates of chromosomal rearrangement are statistically different between the four categories using the Mann-Whitney U test with a Bonferroni correction (supplementary table S5).

### Sequence evolution of aphids, phylloxera, and adelgid

We used syntenic gene pairs between phylloxera and pea aphid and between phylloxera and adelgid from the synteny analysis described in the previous section. The Perl script *add ka and ks to collinearity.pl* from MCScanX (Wang et al. 2012) was used to calculate synonymous (d*S*) and nonsynonymous (d*N*) substitution rates for each syntenic gene pair between the two species. Gene pairs with d*N* > 1 or d*S* > 2 were removed from the analyses to exclude low accuracy estimates of divergence. In phylloxera and pea aphid, we found 226 gene pairs on the X chromosome, and 3,220 gene pairs on autosomes. In phylloxera and adelgid, we found 436 gene pairs on the X chromosome, and 3,758 gene pairs on autosomes (supplementary table S6). We also calculated the d*N*/d*S* ratio for all gene pairs. The mean d*S,* d*N,* and d*N*/d*S* ratios were compared between the X chromosome and autosomes with a Mann-Whitney U test.

### Gene duplication in aphids

Classification of the types of gene duplication was accomplished with the duplicate gene classifier program in MCScanX (Wang et al. 2012). The protein sequences for each species was used as the query and the database in a blastp search with an *e*-value of 1e-10. For each genome, the blastp output and a protein annotation file were used as input files for the duplicate gene classifier program. All default parameters were used. The duplicate gene classifier identified genes as duplications if they hit any other proteins in the blastp search or singletons if they did not. The duplications were further classified into (1) tandem, if they differed by one gene rank; (2) proximal, if they differed by more than 1 and less than 20 gene ranks; (3) dispersed, if they differed by greater than 20 gene ranks; or (4) whole genome duplication (WGD) or segmental duplication, if they were anchored within collinear blocks of genes according to MCScanX (Wang et al. 2012). The percentages for each mode of gene duplication were calculated as the number of duplicate genes in each mode out of the total number of gene duplications in each species (supplementary table S7).

### Identification of putative HGT events

We used Alienness (Rancurel et al. 2017) to identify horizontal gene transfer (HGT) genes in the *D. vitifoliae*. The genome protein sequence file was used as the query against the UniProt Swiss-Prot protein database in a blastp search with an *e*-value of 1e-5 as a cutoff. The blastp output was uploaded to the Alienness website (http://alienness.sophia.inra.fr/cgi/tool.cgi) to identify candidate HGT genes. Metazoa was selected as the group of interest, and *D. vitifoliae* was chosen to be excluded from the calculation of the Alienness Index (AI). Genes with an AI of greater than 30 and percent identity of less than 70% were considered to be candidate HGT genes (Thorpe et al. 2018; Danchin et al. 2017). This filtering process resulted in 194 HGT candidates (supplementary table S8). These protein sequences were checked using blastp against the NCBI non-redundant database with defaults. Proteins with best hits to other arthropod sequences were excluded, resulting in four putative HGT genes.

### PCR amplification of HGT genes

Polymerase chain reaction (PCR) was used to confirm the presence of the four putative HGT genes in the grape phylloxera genome. Phylloxera genomic DNA was extracted from 10 asexual female individuals using the DNeasy Blood & Tissue Kit according to the manufacturer’s protocol for total DNA purification from insects (Qiagen, Germantown, MD, USA). PCR was then used to amplify each of the four HGT genes from the genomic DNA with the primers listed in supplementary table S10. The PCR conditions were as follows: initial denaturation at 95℃ for 30 s; 30 cycles of denaturation at 95℃ for 30 s, annealing at various temperatures (51℃ for plasmid ORF1 and ORF2, 55℃ for glycoside hydrolase 1, 52℃ for glycoside hydrolase 2) for 30 s, and extension at 68℃ for 1 min (plasmid ORFs) or 2 min (glycoside hydrolases); and final extension at 68℃ for 5 minutes. Sanger sequencing was performed by Eton Bioscience, Inc. (San Diego, CA, USA) with the same primers and confirmed the presence of the four HGT genes in the grape phylloxera genome (supplementary fig. S4).

### Transcriptome sequencing and phylogenetic placement of HGT events

To study the four putative HGT genes across aphids, adelgids, and phylloxera, we assembled a dataset of multiple aphid genomes and two adelgid transcriptomes. Given that no other phylloxera transcriptome is available on GenBank besides *Daktulosphaira vitifoliae,* we generated two transcriptomes of *Phylloxera* sp. (one located in galls on hickory leaves and one located in galls on pecan leaves) and a transcriptome of *Phylloxerina nyssae* (BioProject: PRJNA929654, Accession: SRR23289299, SRR23290230, SRR23290233) (supplementary table S11). For each transcriptome, multiple galls were dissected, and insects were pooled into 1.7 mL microcentrifuge tubes. RNA was then extracted from each sample using the Quick-RNA MiniPrep Kit according to the manufacturer’s protocol (Zymo Research, Irvine, CA). The total amount of RNA in each sample was subsequently quantified with a Qubit 2.0 fluorometer (Life Technologies, Carlsbad, CA, USA), and RNA integrity was checked with 4200 TapeStation (Agilent Technologies, Palo Alto, CA, USA). Sequencing libraries were validated with 4200 TapeStation and quantified using a Qubit 2.0 fluorometer and with quantitative PCR (Applied Biosystems, Carlsbad, CA, USA). Sequencing libraries were multiplexed and clustered onto a flow cell and loaded onto an Illumina HiSeq4000 using a 2×150 paired-end (PE) configuration. The sequencing reads were cleaned with Trimmomatic version 0.38 (Bolger et al. 2014). Transcriptomes were assembled using Trinity version v2.1.1 (Grabherr et al. 2011) with default settings.

To assess the presence or absence of the four putative HGT genes, a blastp search with an *e*-value of 1e -2 as a threshold was used to find homologous protein sequences of queries from annotated proteins of genomes and transcriptomes across aphids, adelgids, and phylloxera. The closest hit bacterial protein sequences used as outgroups were also downloaded from Genbank. For each HGT gene family plus outgroup sequences, homologous protein sequences were aligned using MUSCLE v3.8.31 (Edgar 2004) with default parameters. Sequences with low overlap in the protein alignments were removed using -resoverlap 0.75 -seqoverlap 60 in trimAl v1.2 (Capella-Gutiérrez et al. 2009). The ‘-gappyout’ option was used to remove columns with many gaps from the protein sequence alignments. We then manually inspected each protein alignment and removed any ambiguous sequences. Finally, gene trees were built with IQ-TREE multicore version 1.6.1 with 1000 bootstrap replicates using models selected by MFP ModelFinder (Nguyen et al. 2015).

### RNAseq differential expression analyses of HGT genes

Differential gene expression analyses were performed to determine if HGT genes were differentially expressed in the eggs (containing embryos), gallicoles (leaf galls), or radicicoles (root galls) (supplementary table S9). The egg, gallicole, and radicicole transcriptome sequencing reads were obtained from previous studies (Rispe et al. 2016; Zhao et al. 2019) through GenBank (BioProject: PRJNA294954, PRJNA561603). We used the same approach from the previous section for sequence read cleaning. The clean reads were mapped to the chromosome-level assembly using HISAT2 version 2.1.0 (Kim et al. 2019) with -k 3. We used featureCounts (Liao et al. 2014) to estimate the number of reads mapped to the exons of each candidate gene (“–type exon”). Counts of the genes were normalized, and we identified differentially expressed genes using DESeq2 version 1.20.0 in R (Team 2014; Love et al. 2014). The eggs, gallicoles, and radicicoles were each treated as a separate condition in the analysis.

The Wald significance test was used to identify differentially expressed genes. Genes that were >1.5-fold in one condition compared to the other and with an adjusted p-value of 0.05 were identified as significantly affected.

### Data Availability

*Daktulosphaira vitifoliae* chromosome-level genome assembly generated in this study have been submitted to the NCBI BioProject database (https://www.ncbi.nlm.nih.gov/bioproject/) under accession number PRJNA870220. The proximity ligation reads data generated in this study have been submitted to the NCBI BioProject database under accession number PRJNA588186. The sexual male genomic DNA sequencing of the Arizona population generated in this study have been submitted to the NCBI BioProject database under accession number PRJNA929591. The RNAseq sequencing data generated in this study have been submitted to the NCBI BioProject database under accession number PRJNA929654.

## Acknowledgments

This work was supported by the NIH award R35GM131738 to NAM. This project has also received funding from the European Union’s Horizon 2020 research and innovation program under the Marie Sklodowska-Curie grant agreement No 764840. ZL is supported by the National Science Foundation Postdoctoral Research Fellowships in Biology Program (NSF 2109306). We acknowledge Fabrice Legeai and Denis Tagu (INRAE, UMR IGEPP, France), Claude Rispe (INRAE, UMR BIOEPAR, France), and François Delmotte (INRAE, UMR SAVE, France) for discussion and assistance in this study. We thank Eli Powell and Tyler de Jong for assisting with DNA and RNA extractions. We thank Jo-anne Holley and PJ Lariviere for assisting with insect collecting. We also thank Qiuyu Jiang, Michael McKibben, Geoff Finch, and Chenlu Di for collecting the Arizona population of the grape phylloxera.

## References

Blackman RL. 1980. Chromosome numbers in the Aphididae and their taxonomic significance. Syst Entomol. 5:7–25.

Blackman RL, Spence JM. 1996. Ribosomal DNA is frequently concentrated on only one X chromosome in permanently apomictic aphids, but this does not inhibit male determination. Chromosome Res. 4:314–320.

Blackman RL, Spence JM, Normark BB. 2000. High diversity of structurally heterozygous karyotypes and rDNA arrays in parthenogenetic aphids of the genus *Trama* (Aphididae: Lachninae). Heredity 84(2):254–260.

Bolger AM, Lohse M, Usadel B. 2014. Trimmomatic: a flexible trimmer for Illumina sequence data. Bioinformatics 30:2114–2120.

Capella-Gutiérrez S, Silla-Martínez JM, Gabaldón T. 2009. trimAl: a tool for automated alignment trimming in large-scale phylogenetic analyses. Bioinformatics 25:1972–1973.

Chong RA, Park H, Moran NA. 2019. Genome evolution of the obligate endosymbiont *Buchnera aphidicola*. Mol Biol Evol. 36:1481–1489.

Danchin EGJ, Perfus-Barbeoch L, Rancurel C, Thorpe P, Da Rocha M, Bajew S, Neilson R, Guzeeva ES, Da Silva C, Guy J, et al. 2017. The transcriptomes of *Xiphinema index* and *Longidorus elongatus* suggest independent acquisition of some plant parasitism genes by horizontal gene transfer in early-branching nematodes. Genes 8.

Danecek P, Bonfield JK, Liddle J, Marshall J, Ohan V, Pollard MO, Whitwham A, Keane T, McCarthy SA, Davies RM, et al. 2021. Twelve years of SAMtools and BCFtools. Gigascience 10.

Davis GK. 2012. Cyclical parthenogenesis and viviparity in aphids as evolutionary novelties. J Exp Zool B Mol Dev Evol. 318:448–459.

Edgar RC. 2004. MUSCLE: a multiple sequence alignment method with reduced time and space complexity. BMC Bioinform 5:113.

Fernández R, Marcet-Houben M, Legeai F, Richard G, Robin S, Wucher V, Pegueroles C, Gabaldón T, Tagu D. 2020. Selection following gene duplication shapes recent genome evolution in the pea aphid *Acyrthosiphon pisum*. Mol Biol Evol. 37:2601–2615.

Forneck A, Huber L. 2009. (A)sexual reproduction - a review of life cycles of grape phylloxera, *Daktulosphaira vitifoliae*. Entomol Exp Appl. 131:1–10.

Forneck A, Jin Y, Walker A, Blaich R. 1999. Karyotype studies on grape phylloxera (*Daktulosphaira vitifoliae* Fitch). Vitis 38:123–125.

Fradin H, Kiontke K, Zegar C, Gutwein M, Lucas J, Kovtun M, Corcoran DL, Baugh LR, Fitch DHA, Piano F, et al. 2017. Genome architecture and evolution of a unichromosomal asexual nematode. Curr Biol. 27:2928–2939.e6.

Grabherr MG, Haas BJ, Yassour M, Levin JZ, Thompson DA, Amit I, Adiconis X, Fan L, Raychowdhury R, Zeng Q, et al. 2011. Full-length transcriptome assembly from RNA-Seq data without a reference genome. Nat Biotechnol. 29:644–652.

Hardy NB, Peterson DA, von Dohlen CD. 2015. The evolution of life cycle complexity in aphids: ecological optimization or historical constraint? Evolution 69:1423–1432.

Hofstatter PG, Thangavel G, Lux T, Neumann P, Vondrak T, Novak P, Zhang M, Costa L, Castellani M, Scott A, et al. 2022. Repeat-based holocentromeres influence genome architecture and karyotype evolution. Cell 185:3153–3168.e18.

Höök L, Näsvall K, Vila R, Wiklund C, Backström N. 2023. High-density linkage maps and chromosome level genome assemblies unveil direction and frequency of extensive structural rearrangements in wood white butterflies (*Leptidea* spp.). Chromosome Res. 31:2.

Jaquiéry J, Peccoud J, Ouisse T, Legeai F, Prunier-Leterme N, Gouin A, Nouhaud P, Brisson JA, Bickel R, Purandare S, et al. 2018. Disentangling the causes for faster-X evolution in aphids. Genome Biol Evol. 10:507–520.

Jaquiéry J, Rispe C, Roze D, Legeai F, Le Trionnaire G, Stoeckel S, Mieuzet L, Da Silva C, Poulain J, Prunier-Leterme N, et al. 2013. Masculinization of the X chromosome in the pea aphid. PLoS Genet. 9:e1003690.

Jaquiéry J, Stoeckel S, Rispe C, Mieuzet L, Legeai F, Simon J-C. 2012. Accelerated evolution of sex chromosomes in aphids, an X0 system. Mol Biol Evol. 29:837–847.

Johnson KP, Dietrich CH, Friedrich F, Beutel RG, Wipfler B, Peters RS, Allen JM, Petersen M, Donath A, Walden KKO, et al. 2018. Phylogenomics and the evolution of hemipteroid insects. Proc Natl Acad Sci USA 115:12775–12780.

Julca I, Marcet-Houben M, Cruz F, Vargas-Chavez C, Johnston JS, Gómez-Garrido J, Frias L, Corvelo A, Loska D, Cámara F, et al. 2020. Phylogenomics identifies an ancestral burst of gene duplications predating the diversification of Aphidomorpha. Mol Biol Evol. 37:730–756.

Kapustin Y, Souvorov A, Tatusova T, Lipman D. 2008. Splign: algorithms for computing spliced alignments with identification of paralogs. Biol Direct. 3:20.

Kim D, Paggi JM, Park C, Bennett C, Salzberg SL. 2019. Graph-based genome alignment and genotyping with HISAT2 and HISAT-genotype. Nat Biotechnol. 37:907–915.

Kuznetsova VG, Gavrilov-Zimin IA, Grozeva SM, Golub NV. 2021. Comparative analysis of chromosome numbers and sex chromosome systems in Paraneoptera (Insecta). Comp Cytogenet. 15:279–327.

Langmead B, Salzberg SL. 2012. Fast gapped-read alignment with Bowtie 2. Nat Methods 9:357–359.

Li H. 2018. Minimap2: pairwise alignment for nucleotide sequences. Bioinformatics 34:3094–3100.

Li H. 2021. New strategies to improve minimap2 alignment accuracy. Bioinformatics 37:4572–4574.

Li H, Durbin R. 2009. Fast and accurate short read alignment with Burrows-Wheeler transform. Bioinformatics 25:1754–1760.

Li Y, Liu Z, Liu C, Shi Z, Pang L, Chen C, Chen Y, Pan R, Zhou W, Chen X-X, et al. 2022a. HGT is widespread in insects and contributes to male courtship in lepidopterans. Cell 185:2975–2987.e10.

Li Y, Park H, Smith TE, Moran NA. 2019. Gene family evolution in the pea aphid based on chromosome-level genome assembly. Mol Biol Evol. 36:2143–2156.

Li Y, Zhang B, Moran NA. 2020. The aphid X chromosome is a dangerous place for functionally important genes: diverse evolution of hemipteran genomes based on chromosome-level assemblies. Mol Biol Evol. 37:2357–2368.

Li Z, Li Y, Xue AZ, Dang V, Holmes VR, Johnston JS, Barrick JE, Moran NA. 2022b. The genomic basis of evolutionary novelties in a leafhopper. Mol Biol Evol 39.

Liao Y, Smyth GK, Shi W. 2014. featureCounts: an efficient general purpose program for assigning sequence reads to genomic features. Bioinformatics 30:923–930.

Love MI, Huber W, Anders S. 2014. Moderated estimation of fold change and dispersion for RNA-seq data with DESeq2. Genome Biol. 15:550.

Maillet P. 1957. Sur les chromosomes de quelques Phylloxerides de France. Vitis 1:153–155.

Mandrioli M, Manicardi GC. 2020. Holocentric chromosomes. PLoS Genet. 16:e1008918.

Manicardi GC, Mandrioli M, Blackman RL. 2015. The cytogenetic architecture of the aphid genome. Biol Rev Camb Philos Soc. 90:112–125.

Mao M, Yang X, Bennett GM. 2018. Evolution of host support for two ancient bacterial symbionts with differentially degraded genomes in a leafhopper host. Proc Natl Acad Sci USA 115:E11691–E11700.

Mathers TC, Chen Y, Kaithakottil G, Legeai F, Mugford ST, Baa-Puyoulet P, Bretaudeau A, Clavijo B, Colella S, Collin O, et al. 2017. Rapid transcriptional plasticity of duplicated gene clusters enables a clonally reproducing aphid to colonise diverse plant species. Genome Biol. 18:27.

Mathers TC, Mugford ST, Percival-Alwyn L, Chen Y, Kaithakottil G, Swarbreck D, Hogenhout SA, van Oosterhout C. 2019. Sex-specific changes in the aphid DNA methylation landscape. Mol Ecol. 28:4228–4241.

Mathers TC, Wouters RHM, Mugford ST, Swarbreck D, van Oosterhout C, Hogenhout SA. 2021. Chromosome-scale genome assemblies of aphids reveal extensively rearranged autosomes and long-term conservation of the X chromosome. Mol Biol Evol. 38:856–875.

McKenna DD, Shin S, Ahrens D, Balke M, Beza-Beza C, Clarke DJ, Donath A, Escalona HE, Friedrich F, Letsch H, et al. 2019. The evolution and genomic basis of beetle diversity. Proc Natl Acad Sci USA 116:24729–24737.

Miyazaki T, Oba N, Park EY. 2020. Structural insight into the substrate specificity of *Bombyx mori* β-fructofuranosidase belonging to the glycoside hydrolase family 32. Insect Biochem Mol Biol. 127:103494.

Monti V, Lombardo G, Loxdale HD, Manicardi GC, Mandrioli M. 2012a. Continuous occurrence of intra-individual chromosome rearrangements in the peach potato aphid, *Myzus persicae* (Sulzer) (Hemiptera: Aphididae). Genetica 140:93–103.

Monti V, Mandrioli M, Rivi M, Manicardi GC. 2012b. The vanishing clone: karyotypic evidence for extensive intraclonal genetic variation in the peach potato aphid, *Myzus persicae* (Hemiptera: Aphididae). Biol J Linn Soc. 105:350–358.

Moran NA. 1992. The evolution of aphid life cycles. Annu Rev Entomol 37:321–348.

Moran NA. 1988. The evolution of host-plant alternation in aphids: evidence for specialization as a dead end. Am Nat. 132:681–706.

Moran NA, Jarvik T. 2010. Lateral transfer of genes from fungi underlies carotenoid production in aphids. Science 328:624–627.

Moran NA, McLaughlin HJ, Sorek R. 2009. The dynamics and time scale of ongoing genomic erosion in symbiotic bacteria. Science 323:379–382.

Morgan TH. 1909. Sex determination and parthenogenesis in phylloxerans and aphids. Science 29:234–237.

Morgulis A, Gertz EM, Schäffer AA, Agarwala R. 2006. WindowMasker: window-based masker for sequenced genomes. Bioinformatics 22:134–141.

Nakabachi A, Ishida K, Hongoh Y, Ohkuma M, Miyagishima S-Y. 2014. Aphid gene of bacterial origin encodes a protein transported to an obligate endosymbiont. Curr Biol. 24:R640–R641.

Nguyen L-T, Schmidt HA, von Haeseler A, Minh BQ. 2015. IQ-TREE: a fast and effective stochastic algorithm for estimating maximum-likelihood phylogenies. Mol Biol Evol. 32: 268–274.

Normark BB. 1999. Evolution in a putattvely ancient asexual aphid lineage: recombination and rapid karyotype change. Evolution 53:1458–1469.

Nováková E, Moran NA. 2012. Diversification of genes for carotenoid biosynthesis in aphids following an ancient transfer from a fungus. Mol Biol Evol. 29: 313–323.

O’Leary NA, Wright MW, Brister JR, Ciufo S, Haddad D, McVeigh R, Rajput B, Robbertse B, Smith-White B, Ako-Adjei D, et al. 2016. Reference sequence (RefSeq) database at NCBI: current status, taxonomic expansion, and functional annotation. Nucleic Acids Res. 44:D733–45.

Pauchet Y, Heckel DG. 2013. The genome of the mustard leaf beetle encodes two active xylanases originally acquired from bacteria through horizontal gene transfer. Proc Biol Sci. 280:20131021.

Pedersen BS, Quinlan AR. 2018. Mosdepth: quick coverage calculation for genomes and exomes. Bioinformatics 34:867–868.

Purandare SR, Bickel RD, Jaquiery J, Rispe C, Brisson JA. 2014. Accelerated evolution of morph-biased genes in pea aphids. Mol Biol Evol. 31:2073–2083.

Putnam NH, O’Connell BL, Stites JC, Rice BJ, Blanchette M, Calef R, Troll CJ, Fields A, Hartley PD, Sugnet CW, et al. 2016. Chromosome-scale shotgun assembly using an in vitro method for long-range linkage. Genome Res. 26:342–350.

Rancurel C, Legrand L, Danchin EGJ. 2017. Alienness: rapid detection of candidate horizontal gene transfers across the tree of life. Genes 8.

Ren Z, Zhong Y, Kurosu U, Aoki S, Ma E, von Dohlen CD, Wen J. 2013. Historical biogeography of Eastern Asian–Eastern North American disjunct Melaphidina aphids (Hemiptera: Aphididae: Eriosomatinae) on *Rhus* hosts (Anacardiaceae). Mol Phylogenet Evol. 69:1146–1158.

Rispe C, Legeai F, Nabity PD, Fernández R, Arora AK, Baa-Puyoulet P, Banfill CR, Bao L, Barberà M, Bouallègue M, et al. 2020. The genome sequence of the grape phylloxera provides insights into the evolution, adaptation, and invasion routes of an iconic pest. BMC Biol. 18:90.

Rispe C, Legeai F, Papura D, Bretaudeau A, Hudaverdian S, Le Trionnaire G, Tagu D, Jaquiéry J, Delmotte F. 2016. De novo transcriptome assembly of the grapevine phylloxera allows identification of genes differentially expressed between leaf-and root-feeding forms. BMC Genom. 17:219.

Robinson JT, Thorvaldsdottir H, Turner D, Mesirov JP. 2023. igv.js: an embeddable JavaScript implementation of the Integrative Genomics Viewer (IGV). Bioinformatics 39.

Roy SW. 2021. Inbreeding, male viability, and the remarkable evolutionary stability of the aphid X chromosome. Heredity 127:135–140.

Ruckman SN, Jonika MM, Casola C, Blackmon H. 2020. Chromosome number evolves at equal rates in holocentric and monocentric clades. PLoS Genet. 16:e1009076.

Schubert I, Lysak MA. 2011. Interpretation of karyotype evolution should consider chromosome structural constraints. Trends Genet. 27:207–216.

Seppey M, Manni M, Zdobnov EM. 2019. BUSCO: assessing genome assembly and annotation completeness. Methods Mol Biol. 1962:227–245.

Shigenobu S, Watanabe H, Hattori M, Sakaki Y, Ishikawa H. 2000. Genome sequence of the endocellular bacterial symbiont of aphids *Buchnera* sp. APS. Nature 407:81–86.

Simão FA, Waterhouse RM, Ioannidis P, Kriventseva EV, Zdobnov EM. 2015. BUSCO: assessing genome assembly and annotation completeness with single-copy orthologs. Bioinformatics 31:3210–3212.

Smith TE, Li Y, Perreau J, Moran NA. 2022. Elucidation of host and symbiont contributions to peptidoglycan metabolism based on comparative genomics of eight aphid subfamilies and their *Buchnera*. PLoS Genet. 18:e1010195.

So WL, Nong W, Xie Y, Baril T, Ma H-Y, Qu Z, Haimovitz J, Swale T, Gaitan-Espitia JD, Lau KF, et al. 2022. Myriapod genomes reveal ancestral horizontal gene transfer and hormonal gene loss in millipedes. Nat Commun. 13:3010.

Spence JM, Blackman RL. 2000. Inheritance and meiotic behaviour of a de novo chromosome fusion in the aphid *Myzus persicae* (Sulzer). Chromosoma 109:490–497.

The International Aphid Genomics Consortium. 2010. Genome sequence of the pea aphid *Acyrthosiphon pisum*. PLoS Biology 8: e1000313. doi:10.1371/journal.pbio.1000313

Thorpe P, Escudero-Martinez CM, Cock PJA, Eves-van den Akker S, Bos JIB. 2018. Shared transcriptional control and disparate gain and loss of aphid parasitism genes. Genome Biol Evol. 10:2716–2733.

Tree of Sex Consortium. 2014. Tree of Sex: a database of sexual systems. Sci Data 1: 140015.

R Core Team. 2014. R: A language and environment for statistical computing. Vienna: R Foundation for Statistical Computing.

Wang Y, Tang H, Debarry JD, Tan X, Li J, Wang X, Lee T-H, Jin H, Marler B, Guo H, et al. 2012. MCScanX: a toolkit for detection and evolutionary analysis of gene synteny and collinearity. Nucleic Acids Res. 40:e49.

Wickham H. 2016. ggplot2: elegant graphics for data analysis. New York: Springer-Verlag.

Zdobnov EM, Tegenfeldt F, Kuznetsov D, Waterhouse RM, Simão FA, Ioannidis P, Seppey M, Loetscher A, Kriventseva EV. 2017. OrthoDB v9.1: cataloging evolutionary and functional annotations for animal, fungal, plant, archaeal, bacterial and viral orthologs. Nucleic Acids Res. 45:D744–D749.

Zhao C, Nabity PD. 2017. Phylloxerids share ancestral carotenoid biosynthesis genes of fungal origin with aphids and adelgids. PLoS One 12:e0185484.

Zhao C, Rispe C, Nabity PD. 2019. Secretory RING finger proteins function as effectors in a grapevine galling insect. BMC Genom. 20.

