## Supplemental fig. S1-5 for "Phylloxera and aphids show distinct features of genome evolution despite similar reproductive modes"

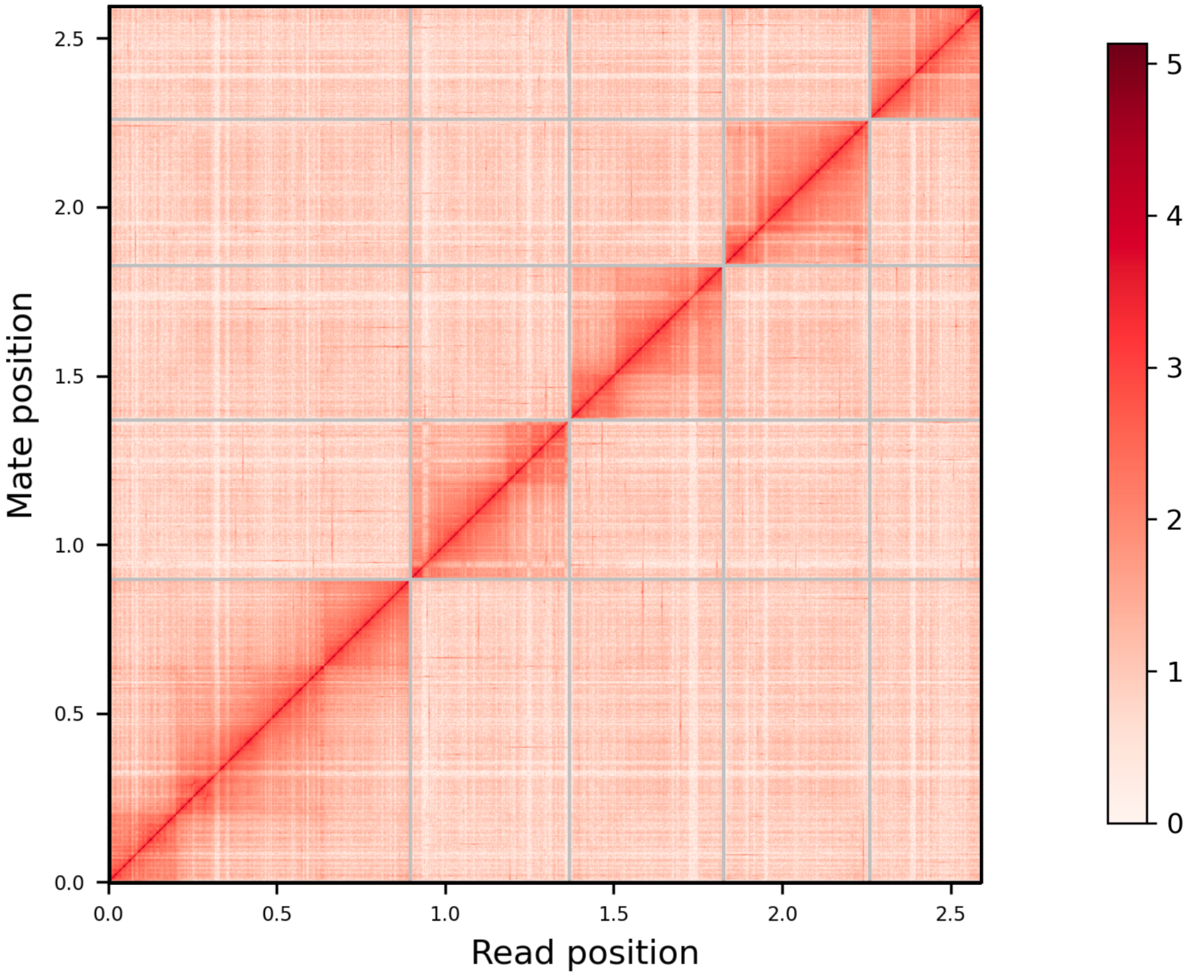


**Supplementary fig. S1** Hi-C contact map of the Dovetail HiRise genome assembly. The X and Y axes indicate the mapping position for the first and second read of each read pair. The color indicates the number of read pairs mapping to each bin. Five chromosome-level scaffolds ordered from largest to smallest.

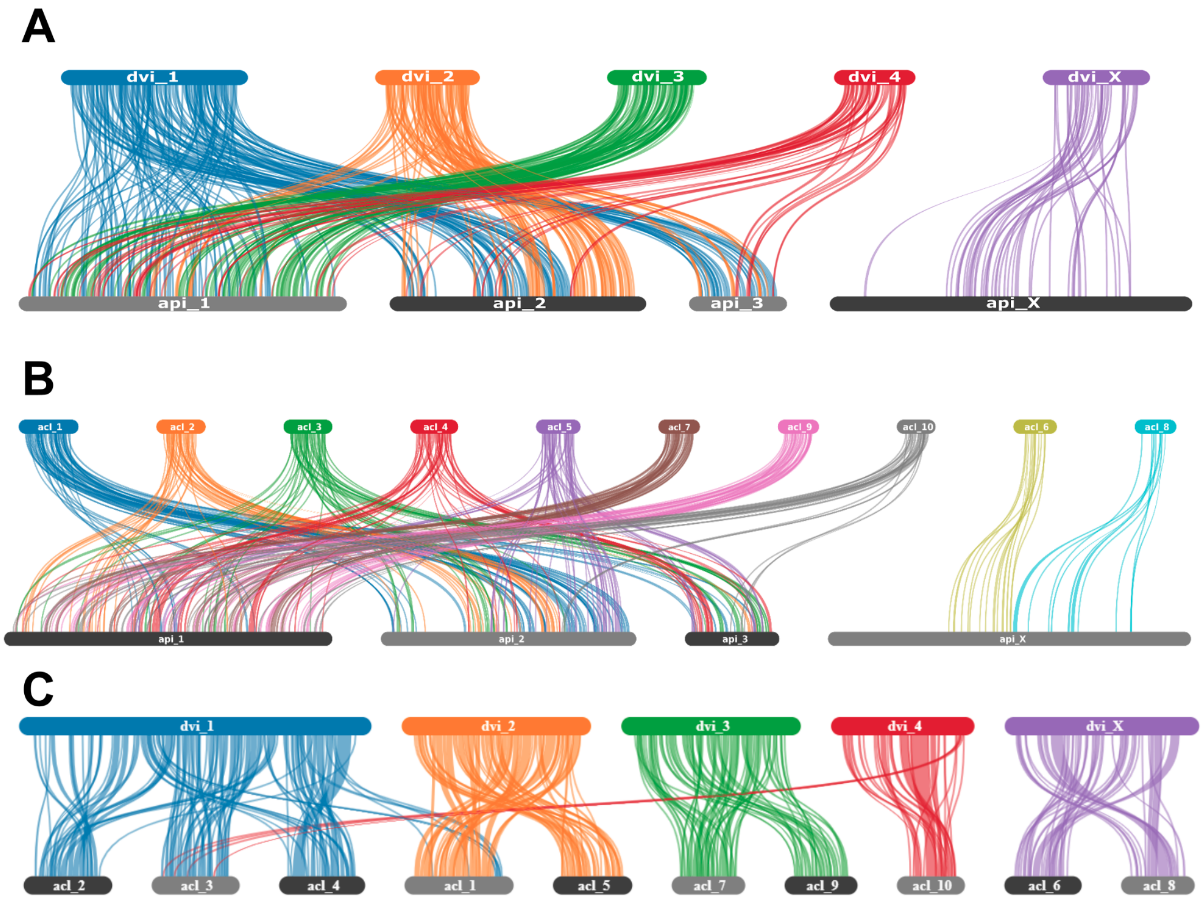


**Supplementary fig. S2** Syntenies (gene order) between **A.** *Daktulosphaira vitifoliae* (dvi, grape phylloxera) and *Acyrthosiphon pisum* (api, pea aphid); **B.** *Adelges cooleyi* (acl, spruce gall adelgid) and *Acyrthosiphon pisum* (api); **C.** *Daktulosphaira vitifoliae* (dvi) and *Adelges cooleyi* (acl). Bars in *Acyrthosiphon pisum* and *Daktulosphaira vitifoliae* represent chromosomes. Bars in *Adelges cooleyi* represent the top 10 largest scaffolds. The length of the bars is proportional to the length of the scaffolds in the assemblies.

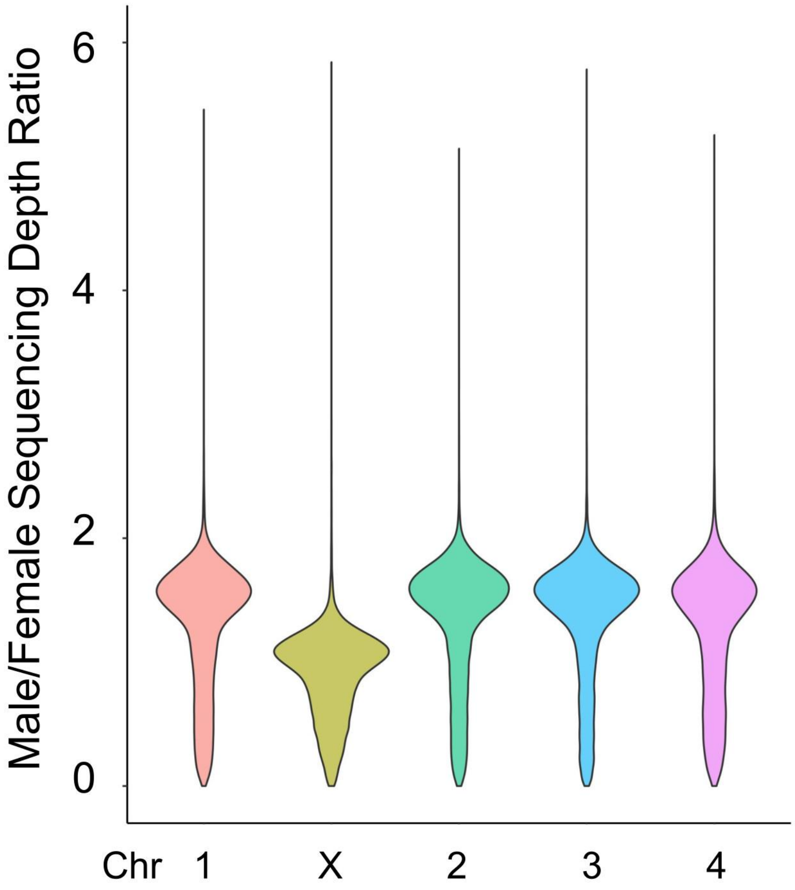


**Supplementary fig. S3** Normalized sequencing depth between male and females of the four autosomes and the X chromosome. The X chromosome showed about half of the sequencing depth ratio between sexes compared to the ratios found for autosomes.

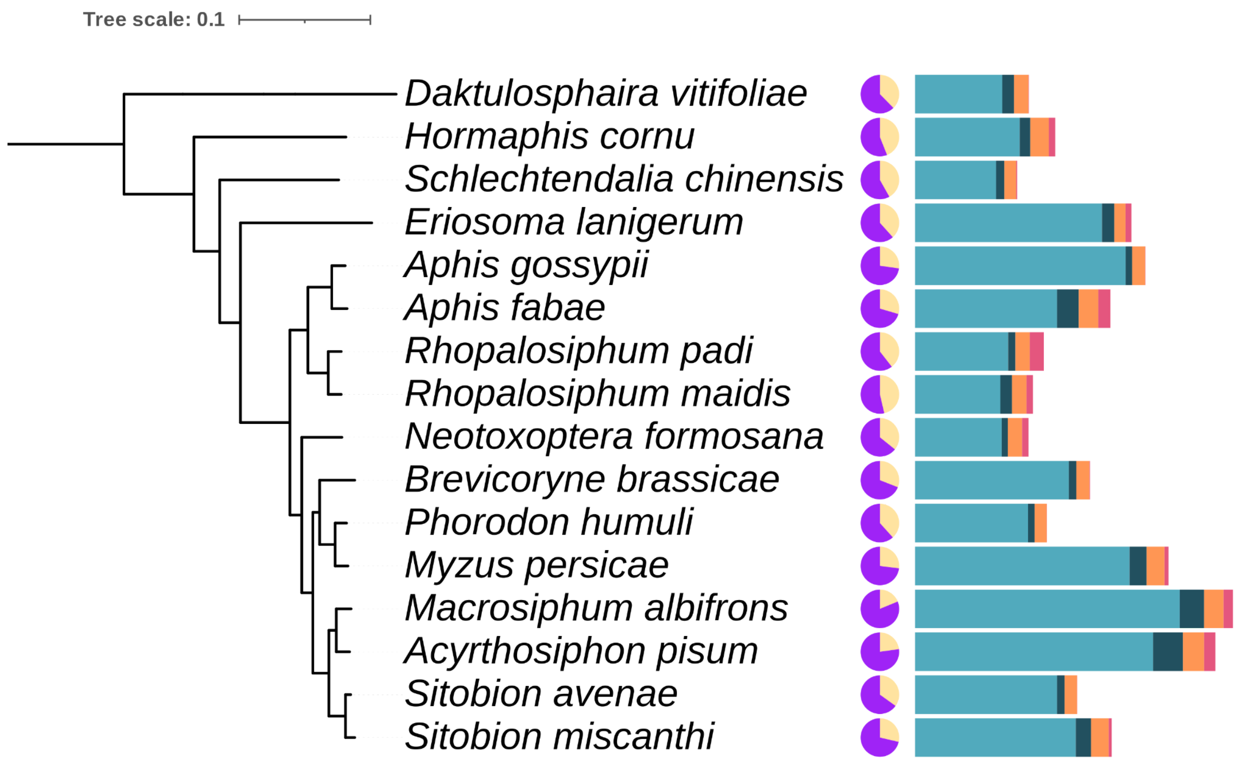


**Supplementary fig. S4** Gene duplications in chromosome-level aphid genomes. The proportion of singletons (yellow) and duplicate genes (purple) for each species are shown in the pie chart. Gene duplications were classified into four different modes: dispersed, proximal, tandem, and segmental duplications (ordered left to right on the bar chart: blue, teal, orange, and red).

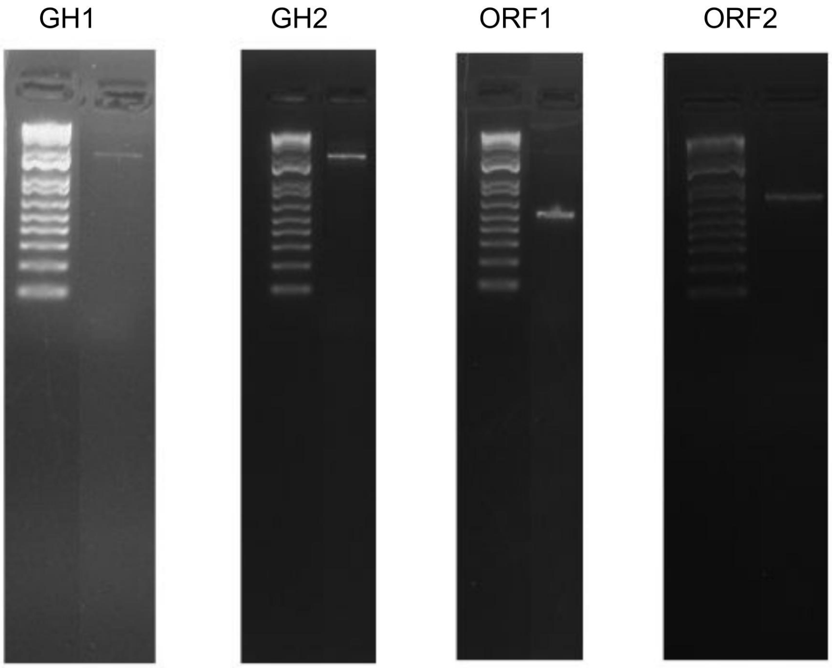


**Supplemental fig. S5** Gel electrophoresis of HGT genes following PCR amplification. Bands represent each of the four putative HGT genes after amplification from phylloxera genomic DNA using PCR, along with 1 kb Plus ladder. From left to right: glycoside hydrolase (GH) 1, GH2, plasmid ORF1, and plasmid ORF2.
